# Typicality of Functional Connectivity robustly captures motion artifacts in rs-fMRI across datasets, atlases and preprocessing pipelines

**DOI:** 10.1101/2020.03.06.980193

**Authors:** Jakub Kopal, Anna Pidnebesna, David Tomeček, Jaroslav Tintěra, Jaroslav Hlinka

## Abstract

Functional connectivity analysis of resting state fMRI data has recently become one of the most common approaches to characterizing individual brain function. It has been widely suggested that the functional connectivity matrix, calculated by correlating signals from regions of interest, is a useful approximate representation of the brain’s connectivity, potentially providing behaviorally or clinically relevant markers. However, functional connectivity estimates are known to be detrimentally affected by various artifacts, including those due to in-scanner head motion. Treatment of such artifacts poses a standing challenge because of their high variability. Moreover, as individual functional connections generally covary only very weakly with head motion estimates, motion influence is difficult to quantify robustly, and prone to be neglected in practice. Although the use of individual estimates of head motion, or group-level correlation of motion and functional connectivity has been suggested, a sufficiently sensitive measure of individual functional connectivity quality has not yet been established. We propose a new intuitive summary index, the Typicality of Functional Connectivity, to capture deviations from normal brain functional connectivity pattern. Based on results of resting state fMRI for 245 healthy subjects we show that this measure is significantly correlated with individual head motion metrics. The results were further robustly reproduced across atlas granularity and preprocessing options, as well as other datasets including 1081 subjects from the Human Connectome Project. The Typicality of Functional Connectivity provides individual proxy measure of motion effect on functional connectivity and is more sensitive to inter-individual variation of motion than individual functional connections. In principle it should be sensitive also to other types of artifacts, processing errors and possibly also brain pathology, allowing wide use in data quality screening and quantification in functional connectivity studies as well as methodological investigations.

## 1. Introduction

Imaging techniques play a pivotal role in medical research nowadays. Functional magnetic resonance imaging (fMRI) is one of the most common methods for research into brain function. Resting-state fMRI (rs-fMRI) is a very prolific and popular subcategory of fMRI measurements. In 1995, Biswal and colleagues found that the correlation of low frequency fluctuations (<≈ 0.1 Hz) in blood oxygen level dependent (BOLD) signal is a manifestation of functional connectivity of the brain. Later studies confirmed that fMRI fluctuations are tightly coupled with the underlying neural activity (Nir et al., 2006; Scholvinck et al., 2010). These spontaneous low-frequency fluctuations in the BOLD signal are therefore used to investigate the functional architecture of the brain (Lee et al., 2013).

A common approach to the analysis of rs-fMRI data is to assess functional connectivity (FC), defined as temporal dependence of neuronal activity patterns (Friston et al., 1993), and thus determine which regions are functionally connected. Region are defined by some reasonable parcellation. Although there is no consensus on optimal brain parcellation (Arslan et al., 2018; Eickhoff et al., 2018), it has been suggested that the matrix of FC among all brain regions may be a suitable representation of the brain connectivity, potentially providing behaviorally or clinically relevant markers (Van Dijk et al., 2009; Biswal et al., 2010; Buckner et al., 2013).

Like any other imaging technique, fMRI is also affected by unwanted artefacts. There are many non-neuronal sources of signal variability such as thermal noise, physiological sources (created by the cardiac and respiratory cycles), scanner and head coil heterogeneities, spiking, chemical shifts, radiofrequency interferences or subject movement (Bianciardi et al., 2009; Chang and Glover, 2009; Poldrack et al., 2011; Murphy et al., 2013). Scanner head motion has long been recognized as a source of artefacts in rs-fMRI (Friston et al., 1996; Hajnal et al., 1994). These artefacts originate in changes in head position that can yield many forms from small involuntary drifts to brief impulsive movements (Patel et al., 2014). They induce undesirable, artificial effects that manifest in complex temporal and spatial patterns (Biswal et al., 1995; Friston et al., 1996; Hajnal et al., 1994; Hlinka et al., 2010; Patel et al., 2014; Spisak et al., 2014). Recent studies showed that even small head movements, in the range of 0.5 to 1 mm, can induce systematic biases in correlation strength and thus they can highly influence the final estimates of functional connectivity (Hlinka et al., 2010; Van Dijk et al., 2012; Power et al., 2012; Satterthwaite et al., 2012; Bright and Murphy, 2013; Mowinckel et al., 2012; Satterthwaite et al., 2013; Tyszka et al., 2014; Yan et al., 2013a). Typical motion artefact manifests as increased short-range connectivity and reduced long-range connectivity, although large head motion can also increase long-range connectivity (Power et al., 2012, 2014, 2015; Satterthwaite et al., 2012, 2019). These effects influence the correlation values as well as the derived connectivity measures characterizing the network topology (Yan et al., 2013b; Ciric et al., 2017). Therefore, they have been both a point of concern and controversy for rs-FC investigations (Bright and Murphy, 2013; Carbonell et al., 2011; Shmueli et al., 2007; Muschelli et al., 2014; Fair et al., 2013; Maclaren et al., 2013).

In common practice, fMRI data preprocessing is used to reduce noise. Preprocessing usually includes image re-alignment, spatial smoothing, filtering, and confound regression (Van Dijk et al., 2009). There is no consensus on optimal preprocessing strategy that should be applied to rs-fMRI data (Aurich et al., 2015). Since no preprocessing is completely successful in removing the motion artefact (Ciric et al., 2017; Siegel et al., 2017) it is vital for connectivity studies to be able to quantify the amount of motion artefacts present in FC matrices. However, a reliable measure of FC quality has not yet been established. Absence of robust FC quality measure renders the estimation of the amount of motion artefact in a FC matrix impossible. We propose a new measure - Typicality of Functional Connectivity, that is based on a correlation of an individual FC matrix with a typical FC matrix. Such measure can be helpful in investigations of individuals and populations whose in-scanner movement profiles may differ subtly, for instance when comparing controls to subjects of different ages (e.g. during development or aging) or to individuals experiencing involuntary or repetitive movements (e.g. tics or tremors) (Muschelli et al., 2014; Bright and Murphy, 2013). By construction, it should be sensitive also to other types of artifacts, processing errors and possibly also brain pathology, allowing wide use in data quality screening and quantification in functional connectivity studies as well as methodological investigations, such as the evaluation of preprocessing pipeline performances and the decision on suitable brain parcellation.

## 2. Material and Methods

### 2.1. Data acquisition

245 healthy subjects (148 right-handed, 132 females, mean age 29.22 / standard deviation 6.99) participated in the study. Participants were informed about the experimental procedures and provided written informed consent. The study design was approved by the local Ethics Committee of the Institute of Clinical and Experimental Medicine and the Psychiatric Center Prague. Each volunteer underwent MRI scanning that included 10 minutes of resting-state functional magnetic resonance imaging acquisitions with eyes closed and an acquisition of a T1-weighted and T2-weighted anatomical scan.

Scanning was performed with a 3T MR scanner (Siemens; Magnetom Trio) located at the Institute of Clinical and Experimental Medicine in Prague, Czech Republic. Functional images were obtained using T2-weighted echo-planar imaging (EPI) with blood oxygenation level-dependent (BOLD) contrast. GE-EPIs (TR/TE=2000/30 ms) comprised axial slices acquired continuously in descending order covering the entire cerebrum (voxel size=3 × 3 × 3mm^3^). A three-dimensional high-resolution T1-weighted image (TR/TE/TI = 2300/4.6/900 ms, voxel =1 × 1 × 1mm^3^) covering the entire brain was used for anatomical reference. T2-weighted images were also acquired, but not used in the current study.

### 2.2. Preprocessing

Initial data preprocessing was performed using a combination of SPM12 software package (Wellcome Department of Cognitive Neurology, London, UK), CONN toolbox (McGovern Institute for Brain Research, MIT, USA) running under MATLAB (The Mathworks) and FSL routines (FMRIB Software Library v5.0, Analysis Group, FMRIB, Oxford, UK). CONNs default preprocessing pipeline (defaultMNI) comprises of the following steps: (1) functional realignment and unwarping, (2) slice-timing correction, (3) structural segmentation into white matter and cerebrospinal fluid & structural normalization to the MNI space, (4) functional normalization to the MNI space, (5) outlier detection, and (6) smoothing with 8mm kernel size.

The denoising steps included regression of six head-motion parameters (acquired while performing the correction of head-motion) with their first-order temporal derivatives and five principal components of white-matter and cerebrospinal fluid. The CONN toolbox has implemented a component-based noise correction method (CompCor) that typically performs PCA dimensionality reduction of white-matter a cerebrospinal fluid time-series derived from specific regions (Behzadi et al., 2007). The CompCor method uses noise regions of interest acquired while segmenting each subjects high-resolution anatomical images (Chai et al., 2012). Time series from defined regions of interest were additionally linearly detrended to remove possible signal drift and finally filtered by a band-pass filter with cutoff frequencies 0.009 - 0.08 Hz. This preprocessing pipeline is labeled as *stringent* further in the manuscript.

To form functional connectivity matrices, we cross-correlated the ROI-based average BOLD time series. In line with the most common practice, we use Pearson correlation coefficient to quantify functional connectivity and form FC matrices. Note that although other nonlinear approaches for functional connectivity assessment have been proposed, linear Pearson correlation coefficient was shown to be sufficient under standard conditions (Hlinka et al., 2011; Hartman et al., 2011). Fisher’s r-to-z transformation (Zar, 1999) was applied to each correlation coefficient to increase normality of the distribution of correlation values.

#### 2.2.1. Atlas choice

A common approach to extract BOLD time series is to use brain parcellations. Brain parcellations divide the brains spatial domain into a set of non-overlapping regions of interest or modules that show some homogeneity with respect to information provided by one or several image modalities, such as cytoarchitecture, task-based fMRI activations, or anatomic delineations (Thirion et al., 2014; Shen et al., 2013).

In our analysis we chose a parcellation based on Craddock atlas because it offers an option to select a number of ROIs that represent spatially coherent regions with homogeneous connectivity. We calculated 23 FC matrices for each subject that were based on atlases of various number of ROIs; ranging from 10 to 840 ROIs. With increasing number of ROIs, the size of each ROI decreases (Fig.1). The regions in Craddock atlas are created using a spectral clustering algorithm with various similarity metrics and group-level clustering schemes (for details see Craddock et al., 2012).

**Figure 1:**
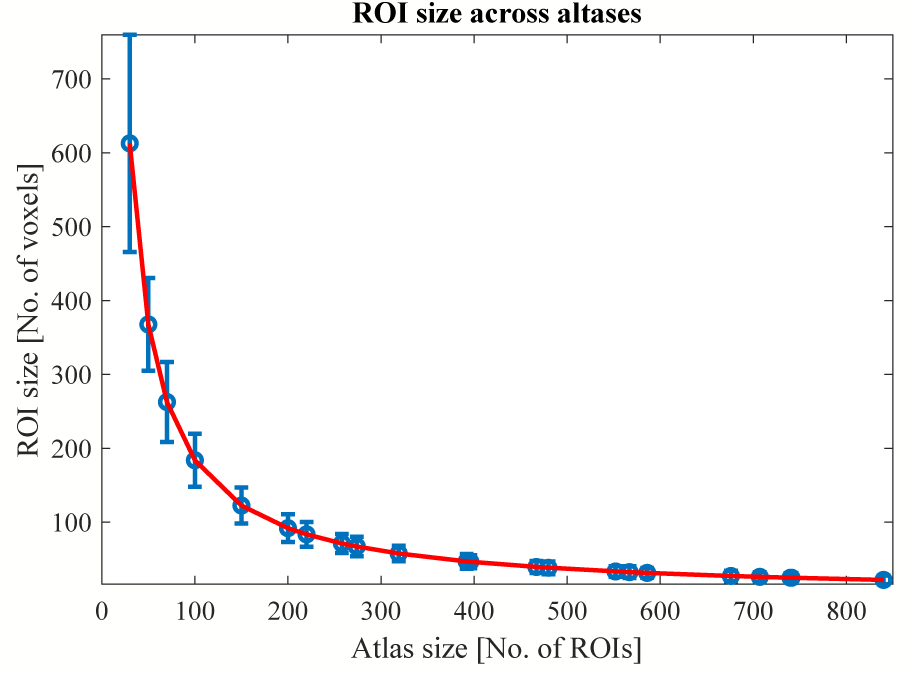
ROI sizes for 23 atlases based on Craddock spectral clustering method. Mean size (number of voxels) with ± standard deviation is plotted. The ROI size decreases with increasing number of ROIs.

### 2.3. Quantifying motion

Reporting motion statistics should be fundamental for any fMRI study but Waheed et al. (2016) analyzed 100 most recent fMRI studies and only 10 % provided a table of motion metrics. Two of the most used motion metrics are framewise displacement (FD) and derivative of root mean square variance over voxels (DVARS). We used mean FD and mean DVARS to quantify the amount of motion during a given scanning session. Distribution of each metric is available in the appendix (Appendix C).

#### 2.3.1. Framewise displacement (FD)

The fMRI data allow estimation of six head realignment parameters for each volume. Thus, head position is described at each time point by six parameters (translational displacements along X, Y, and Z axes and rotational displacements of pitch, yaw, and roll). Framewise displacement represents a summarizing parameter of head motion from one volume to the next. It is an average of the rotation and translation parameters differences (Eq.1). Since it is based on realignment parameters, it is therefore unaffected by subsequent preprocessing steps (Power et al., 2012).

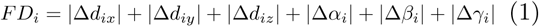

where displacement of *i*-th brain volume in x-direction is Δ*d*_*ix*_ = *d*_(*i*−1)*x*_ − *d*_*ix*_ and similarly for the other rigid body parameters.

Rotational displacements were converted from degrees to millimeters by calculating displacement on the surface of a sphere of radius 50 mm.

FD is the most popular metric among motion statistics. It was reported in 24 % of recent fMRI studies compared to similar root mean square (RMS) metric which was reported only in 10 % of recent fMRI studies (Waheed et al., 2016).

#### 2.3.2. Derivative of root mean square variance over voxels (DVARS)

Derivative of root mean square variance over voxels is based on the fact that abrupt head displacement typically manifests as signal loss in echo-planar imaging. It quantifies changes of intensities between two images and it is calculated as the root mean square value of the differentiated BOLD time series within a spatial mask at every time-point (Eq.2; Smyser et al. (2010)). DVARS is not derived from realignment parameters and thus it can reflect any kind of bias. Nevertheless, head motion has been proven to be a major contributor to fluctuations in DVARS (Fair et al., 2013). The quantity is defined as:

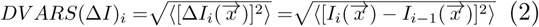

where 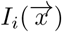 is image intensity at locus 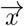 on frame *i* and angle brackets denote the spatial average over the whole brain.

Since it is based on BOLD signal intensity, DVARS differs greatly across datasets and processing strategies (Power et al., 2014). It can be influenced by blurring kernel size, frequency filter characteristics, sequence characteristics, etc. DVARS was reported only in 8 % of the recent fMRI studies (Waheed et al., 2016).

### 2.4. Measuring FC quality

Estimating connectivity quality and assessing its relationship with motion is important for all connectivity studies. Currently there is no measure used in literature that allows quantifying it per subject.

#### 2.4.1. Quality control-functional connectivity (QC-FC)

In literature the most used way to evaluate presence of a motion artefact are quality control-functional connectivity (QC-FC) values (Power et al., 2014; Satterthwaite et al., 2012; Ciric et al., 2017; Parkes et al., 2018). This group measure focuses on influence of motion on connectivity values across subjects. It describes how motion affects FC values. Each correlation coefficient in a FC matrix is directly correlated with a summary motion statistic across subjects. The median of these values shows if motion tends to increase or decrease connectivity and a correlation of QC-FC with distance reveals presence of spurious distance dependence.

#### 2.4.2. Typicality of Functional Connectivity (TFC)

We propose Typicality of Functional Connectivity as a new measure for how to estimate FC quality. It is based on a correlation between an individual subjects FC matrix and a typical FC matrix of a given cohort (Eq.3). To exclude influence of the diagonal, we vectorized upper triangular form of all FC matrices (ignoring the diagonal elements).

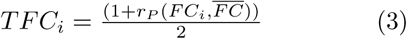

where *i* is the subject’s index, *r*_*P*_ is a Pearson correlation coefficient and 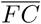 is the typical FC matrix. Throughout the manuscript Spearman correlation is denoted as *r*_*S*_ and Pearson correlation as *r*_*P*_.

TFC ranges between 0 and 1, where 0 is a complete anti-correlation, 0.5 is correlation of 0 and 1 is maximal correlation with the typical FC matrix.

As the template, we use the mean FC matrix of 20 % subjects with lowest motion (49 subjects with lowest mean FD) as the typical matrix - a golden standard, to which other subjects are compared; although taking mean FC across the whole dataset or from a different dataset gives similar results. We assume that by averaging FC matrices of low-movement subjects we obtain a FC matrix representing typical awake human brain functional connectivity free of motion artefact (as well as other biases). While minor or moderate deviations may in principle represent effects of interest corresponding to inter-individual variation in brain function, larger anomalies are likely to be due to artifactual sources of signal variation and should be subject to screening.

## 3. Results

We used TFC to estimate per subject quality and we analyzed it with respect to motion, atlas size and preprocessing. Using Spearman correlation we found that it is significantly correlated with motion metrics 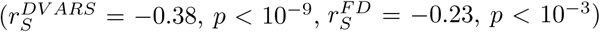, meaning that a FC matrix of a subject with high mean head movement is less similar to the typical FC matrix compared to low-movement subjects (Fig.2a). Such correlation coefficient between a motion metric and TFC metric demonstrates a dependence between FC quality and gross head motion. Both FD and DVARS are significantly related to FC quality but mean DVARS shows generally higher absolute correlation than mean FD. This is most likely due to the fact that DVARS captures other artefacts and impurities as well.

**Figure 2:**
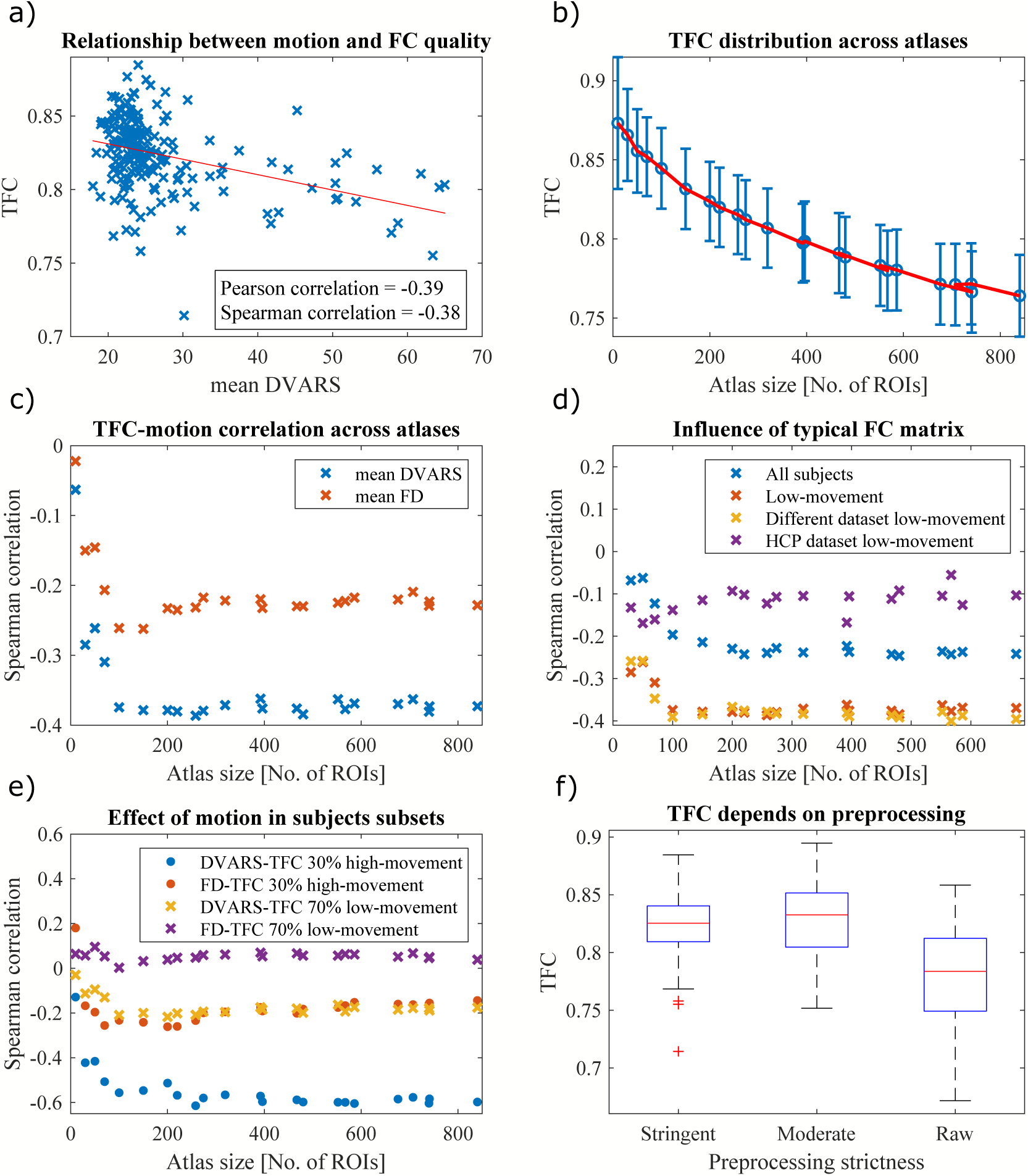
Analysis of new FC quality metric - TFC. a) Relationship between mean DVARS and TFC. Subjects with worse FC quality (lower correlation coefficient between an individual FC matrix and the typical FC matrix) show higher amount of motion during the scanning session. Calculated for Craddock atlas with 200 ROIs. b) Mean ± standard deviation of TFC across atlases. The quality of functional connectivity is decreasing as the number of ROIs increases. c) Spearman correlations between TFC and two summarizing motion metrics for atlases with different number of ROIs. Except for the very small atlases the relationship between FC quality and motion is constant. c) The highest absolute correlation of the TFC-motion dependence is obtained if low-movement subjects of the same dataset are used for the calculation of the typical FC matrix compared to using all subjects or low-motion subjects of different datasets. Such typical matrix is comparable to a typical matrix of a different dataset (*r*_*P*_ = 0.92, *p* < 10^−16^) and similar to a typical matrix of HCP dataset (*r*_*P*_ = 0.68, *p* < 10^−16^). d) Subjects were divided into two subsets - 30 % of subjects with the highest motion and 70 % with the lowest motion. High-movement cohort has a stronger relationship between FC quality and motion compared to low-movement cohort for both DVARS and FD. Low-movement cohort did not display any kind of influence of motion on quality as all the correlation were statistically insignificant. e) Comparison of quality of FC matrices of all subjects for three different preprocessing pipelines with different level of strictness; stringent being the most strict and raw the most lenient. FC matrices with more strict preprocessing are more similar to the typical FC matrix. Moreover, the standard deviation is reduced in more strict preprocessing pipeline.

Instead of TFC, we also tried a method based on Euclidean *L*^2^ distance (Ponsoda et al., 2017) from the typical FC matrix and mean geodesic distance from the cohort (Venkatesh et al., 2020). Unlike TFC measure, which shows significant both Spearman and Pearson quality-motion correlations, the correlations of *L*^2^ distance with motion were significant only for Pearson correlation 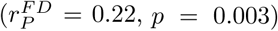 because this relationship was driven mainly by outliers. Correlations with geodesic distance did also yield only one outlier-driven significant result 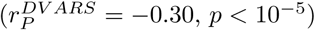 (Tab.1).

**Table 1:** Correlation of different measures of FC quality with motion metrics. Only TFC shows significant correlations for both motion metrics and for both Pearson and Spearman correlations. Relationships with only Pearson correlations significant were driven mainly by outliers. * signifies *p* < 0.05 and ** signifies *p* < 0.01.

|  | DVARs |  | FD |  |
| --- | --- | --- | --- | --- |
|  | Spearman | Pearson | Spearman | Pearson |
| TFC | **-0.38\*\*** | **-0.39\*\*** | **-0.23\*\*** | **-0.26\*\*** |
| L2-distance | 0.05 | -0.02 | 0.03 | **0.22\*\*** |
| Geodesic distance | 0.00 | **-0.30\*\*** | 0.05 | 0.03 |

Since Spearman correlation is less sensitive to outliers compared to Pearson correlation, we prefer to use it throughout the manuscript when assessing the relationship with motion.

We further analyzed only TFC as a quality measure. We evaluated it for every subject across Craddock atlases with varying number and size of ROIs. From Fig.2b it is evident that FC quality decreases as the atlas size decreases. Therefore, more detailed FC matrices are of worst quality. Furthermore, we investigated whether this gradual decrease is driven by increased effect of motion on signal in small regions. We calculated correlations between motion and TFC across variously detailed atlases and found that, except for atlases with less than 100 regions, the relationship is stable (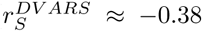 and 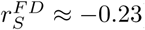) (Fig.2c).

The relationship is the strongest if the typical FC matrix is based on 20 % of subjects with the lowest mean FD of the same dataset and only slightly weaker if different dataset with identical preprocessing pipeline is used (more information about alternative datasets in Appendix A). If all subjects are used for the calculation of the typical FC matrix the observed relationships are weaker (Fig.2d), possibly due to presence of various types of noises. Nevertheless, even using different dataset with different preprocessing, such as Human Connectome Project (HCP, see Appendix F), still gives significant (only for DVARS) results.

To further investigate the effects of head movements we split the dataset into two parts - 30 % of subjects with the highest motion (mean *FD* > 0.2) and 70 % subjects with the lowest motion (mean *FD* < 0.2). Low-movement subjects show only weak and statistically insignificant quality-head movement correlations 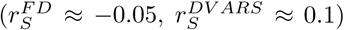 proving that the effect is more pronounced in subjects with higher motion (same effect observed in HCP dataset). These subjects exhibit gradual increase of the dependence for the smallest atlases (10, 30, 50, 70 ROIs), from 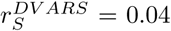 for 10 ROI atlas up to 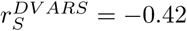 for 100 ROI atlas (Fig.2e). For more detailed atlases (more than 100 ROIs) the relationship is constant. We obtained similar results if the subsets were created based on thresholding DVARS.

Besides the influence of ROI size on FC quality, we also analyzed the influence of data preprocessing on FC quality. We compared FC quality for three different preprocessing pipelines based on their strictness - stringent, moderate and raw (more about pipeline differences in Appendix B). In Fig.2f we see that for the two strict preprocessing pipelines individual FC matrices more resemble the typical FC matrix: mean(*TFC*_*stringent*_) = 0.82, mean(*TFC*_*moderate*_) = 0.83, mean(*TFC*_*raw*_) = 0.78. Standard deviation of quality measure is increasing with decreasing strictness of preprocessing: std(*TFC*_*stringent*_) = 0.02, std(*TFC*_*moderate*_) = 0.03, std(*TFC*_*raw*_) = 0.04. For all these cases we used the typical FC matrix of a dataset with stringent preprocessing, but results were similar if each preprocessing stream used its own FC matrix as a golden standard.

In the literature the most used metric to assess presence of motion artefacts are QC-FC values. Instead of examining how motion affects each subject’s FC matrix, it examines how motion affects FC values for each pair of regions. We obtained positive median of QC-FC and significant negative correlation between QC-FC and distance for both quality metrics 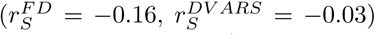 (Fig.3a,b). As reported previously (Power et al., 2012, 2015; Satterthwaite et al., 2019) we confirm that motion affects distance-dependence and on average causes spurious increase in connectivity. Moreover, this effect is constant across atlases of various size (Fig.3d). Nevertheless, only 15 % (resp. 4 %) of QC-FC values were significant (Fig.3c). The main disadvantage of QC-FC values is that it can be used only on a group level and it does not allow single subject description.

**Figure 3:**
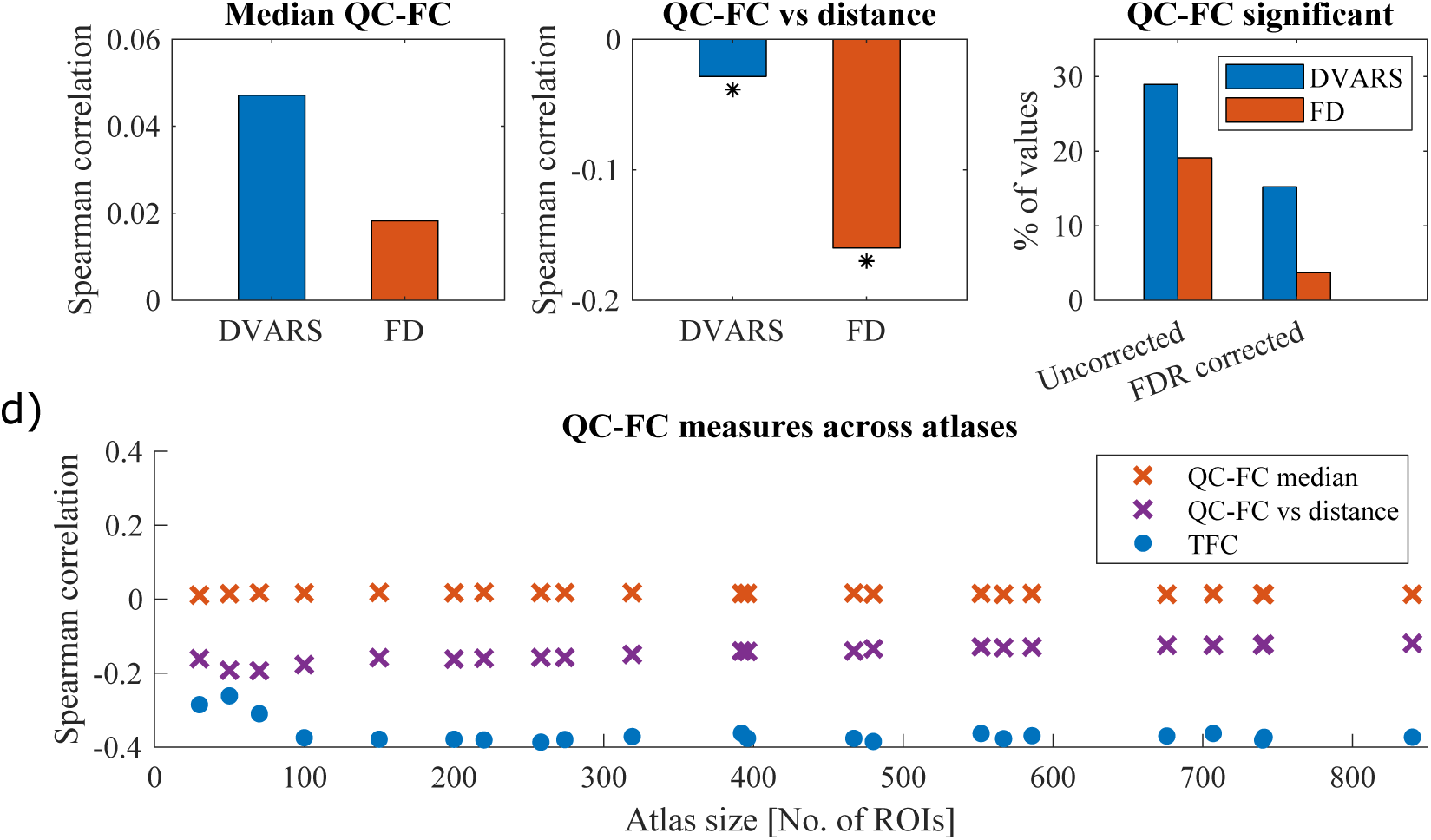
The QCFC correlations quantify the association between inter-individual variance in functional connectivity and gross head motion. a) Positive median of QC-FC values signifies that head motion increases connectivity. b) This effect is more prominent for short-links and it is more specifically related to motion as FD correlations are stronger than DVARS correlations. * signifies *p* < 0.05 c) The amount of edges that is significantly affected by movements. The effect is more easily detectable with DVARS metric. Results are plotted for Craddock atlas with 200 ROIs. d) Above mentioned effects are stable across atlases with different number of ROIs. Magnitudes of TFC correlations are higher than median of QC-FC, proving its viability as an estimator. Plotted only for mean FD.

So far, we focused only on quality of connectivity matrices, but noisiness of underlying BOLD time series can be estimated as well. BOLD signal quality is usually expressed in the form of temporal signal to noise ratio (tSNR). tSNR compares the level of a desired signal to the level of a background noise (for details on calculation see Appendix E). Noise is generally more present in more lenient preprocessing pipelines (Fig.4d). Moreover, we can observe a gradual decrease of tSNR with decreasing ROI size (Fig.4a). To test whether such degradation is related to motion, we correlated FD and DVARS with tSNR across atlases (Fig.4c). DVARS displays progressive increase of an absolute correlation with tSNR unlike FD (change of correlation between smallest and highest atlas: 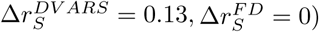). It is apparent that tSNR measures different data aspects compared to TFC as they correlate only weakly (Fig.4b). In conclusion, there is an effect of ROI size on data quality, but it might be more specifically related to other types of noise than a head movement, because only the tSNR-DVARS dependence varied and DVARS captures other impurities as well.

**Figure 4:**
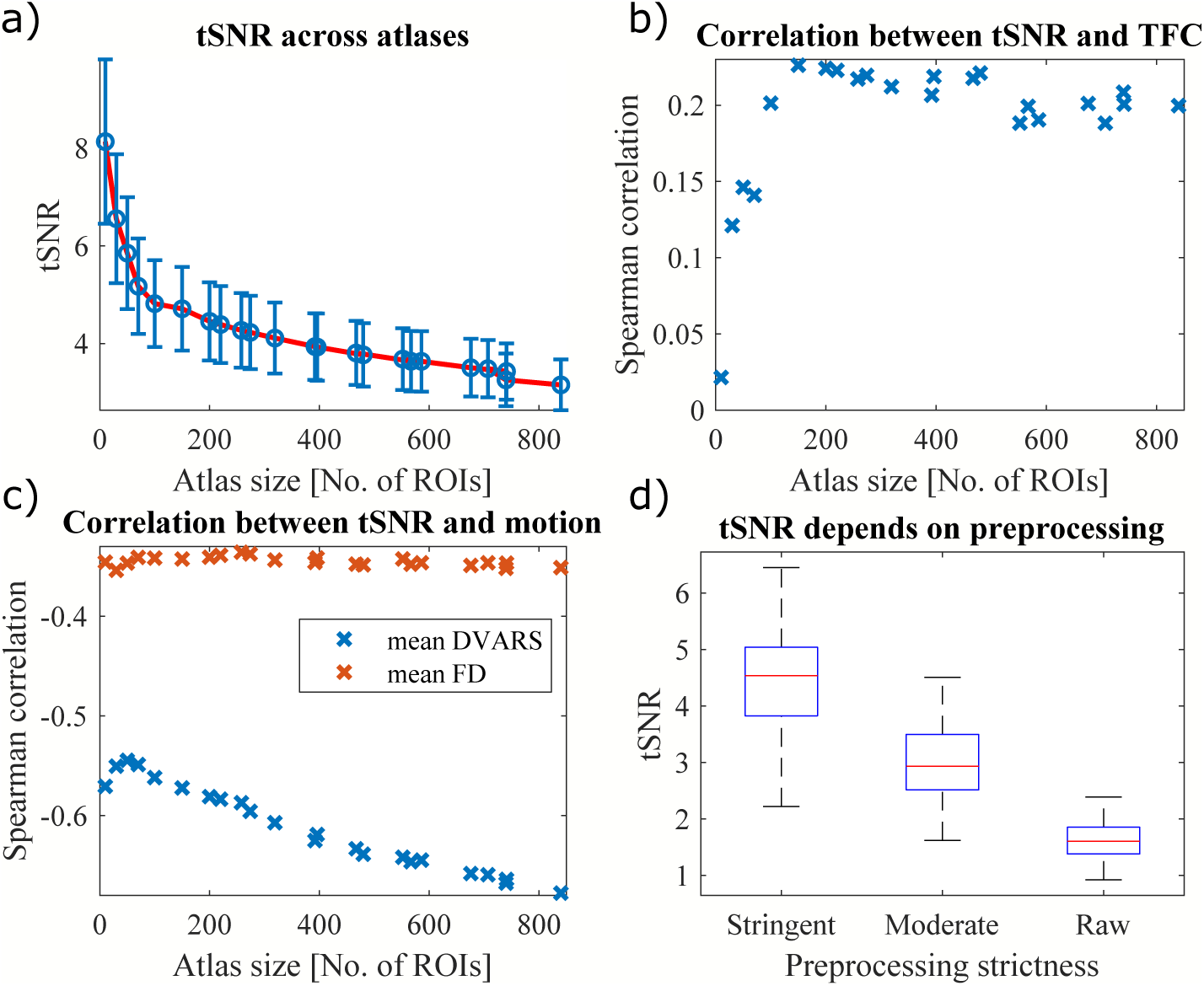
Analysis of tSNR with respect to atlas size, motion and FC quality. a) Mean tSNR ± standard deviation across different atlases. tSNR is gradually decreasing for more detailed atlases, therefore smaller ROIs are more affected by noise. b) tSNR measures different data aspects than TFC as the correlation is weak, but significant and positive. c) With decreasing size of ROIs, the relationship between tSNR and mean DVARS get stronger. This trend is not present for FD meaning that the phenomenon is predominantly caused by other types of noise than a head movement. d) As well as TFC, tSNR also depends on preprocessing, where the more stringent pipeline produces less noisy time-series.

Several times we observed the effect of atlas size for atlas sizes up to 100 ROIs. This effect might be driven by two factors: by the number of regions or by the size of regions. To test the first hypothesis, we randomly selected 50,100,150,…,700 ROIs out of an atlas with 950 ROIs and we calculated both TFC and tSNR and analyzed their relationship with head movement. Such procedure was repeated 1000 times. In such scenario number of voxels in a region is fixed - 21.9 ± 0.3 and only number of regions varies. Neither tSNR nor TFC depends on the number of regions. We only observed small gradual increase in the TFC-motion relationship when only few regions were selected. To test the second hypothesis, we took atlas with 100 ROIs (183.8 ± 35.8 voxels per region) and we created different regions around the central voxel that varied in the number of voxels (Fig.5c). Both tSNR and TFC increase with increasing number of voxels. On the contrary, the TFC-motion dependence is weaker for low number of voxels (Fig.5b). These results suggest that regions with less voxels produced noisier data. Additionally, when choosing only few regions (< 100) it is more difficult to reliably estimate significant relationship between quality and movement.

**Figure 5:**
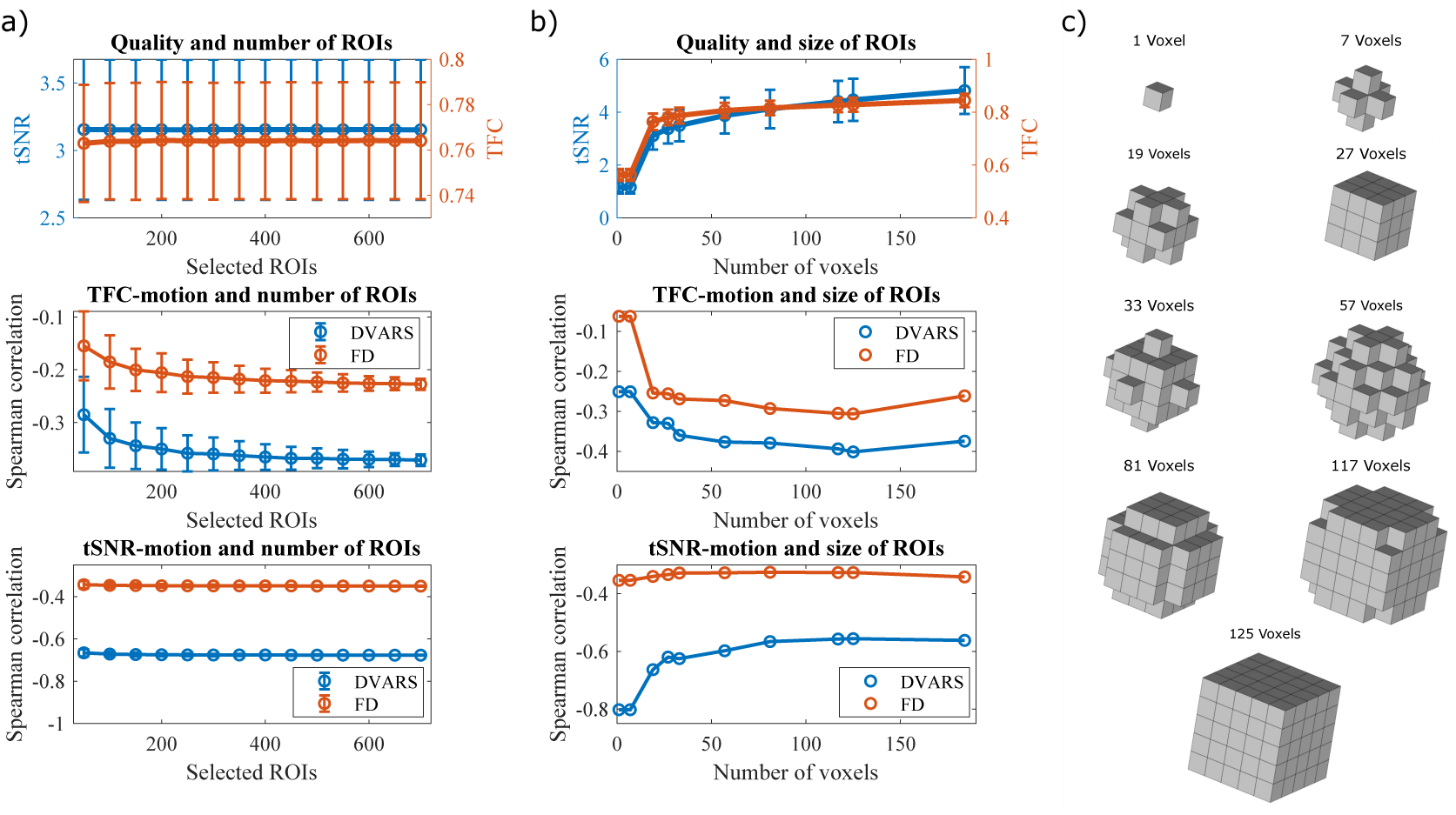
Are atlas size effects driven by number of regions of number of voxels? a) In an atlas with 950 ROIs we randomly selected 50,100,150,…,700 ROIs to get quality estimates depending only on a number of regions but independent of a number of voxels. Only relationship of TFC and motion is slightly weaker for smaller numbers of regions. b) Within an atlas of 100 ROIs we varied the number of voxels that forms a region. Both tSNR and TFC depend on number of voxels. Moreover, while the tSNR-DVARS relationship is stronger for smaller number of voxels, opposite effect is present for the TFC-motion relationships. c) Different geometrical shapes build around a central voxel of a region. Thus, we could vary number of voxels forming a region.

## 4. Discussion

### 4.1. Estimation of FC quality

The lack of gold standard for FC quality estimation has hampered direct comparison among different groups (neurodevelopmental, aging, neuropsychiatric), preprocessing pipelines and brain parcellations. We introduced a new measure (TFC) to describe the quality of a functional connectivity per subject. This measure is based on a correlation of an individual FC matrix with the low-motion group-average connectivity matrix. As we showed, it provides a reliable estimate of FC quality with respect to motion and atlas size and possibly other types of noise. Low-movement subject’s FC matrices are strongly correlated with the typical FC matrix compared to high-movement subjects, despite the fact that even our high-movement subjects were healthy controls and would meet inclusion criteria for analysis in most MRI laboratories (C.6). Moreover, even by visual inspection it is apparent that subjects with low TFC have lost modular structure of the typical FC matrix or they show general artifactual increase in connectivity (D.7).

An alternative measure to TFC could be *L*^2^ distance from the typical FC matrix or mean of geodesic distance from the cohort but our results suggest that these measures are less specifically related to motion. One of the reasons could be that they are more sensitive to other global artefacts.

Currently many studies propose QC-FC values as a measure of motion impact (Power et al., 2015; Ciric et al., 2017; Power et al., 2017). QC-FC values are correlations between vectors of summary quality (motion) control values (e.g., mean FD, mean DVARS, etc.) with vectors of outcome measures (FC values) across subjects. A limitation of this measure is the lack of single subject quality estimate. At a group level they describe motion artefact manifestations. We confirmed that head movements generally increase connectivity (median QC-FC similar to the one reported in Ciric et al. (2017) and Parkes et al. (2018) for corresponding preprocessing pipeline) and that it affects distance dependency - increased short-range connectivity and decreased long-range connectivity. This spatial pattern is specifically related to motion as we found stronger dependence for FD. As reported in Ciric et al. (2017) number of links related to motion varies significantly (in our results more than 80% QC-FC values insignificant). In addition, it could be difficult to reliably establish a QC-FC correlation if there is little variability in the QC measure (Power et al., 2015). Moreover, such measures are sensitive to outlying values and a few scans with marked abnormalities can obscure relationships present across most other datasets Power et al. (2017). That is why QC-FC should be complemented with other assessments.

Several other metrics have also been used as well in prior reports, including FD-DVARS correlations. DVARS was used as a predictor of data quality rather than an estimate of amount of motion. Before the preprocessing, DVARS strongly resembles FD, but this similarity diminishes with processing (Power et al., 2014) and that is why DVARS could serve as a marker of nuisances in a FC matrix (Hallquist et al., 2013; Power et al., 2012, 2014). Nevertheless, DVARS changes during processing steps even when the motion artefact is not filtered out (Spisak et al., 2014), therefore the FD-DVARS relationship is not recommended to estimate FC quality but rather it is advised to use DVARS as a motion metric. Another metric sometimes used to assess the presence of motion artefact and the success of denoising strategies are FD-BOLD signal correlations. It has been suggested that the FD-BOLD correlations reveal motion-related neural activity (Yan et al., 2013a,b). However, according to Power et al. (2015) these correlations are probably not related to neural activity. Finally, Saad et al. (2013) proposed global correlation (i.e. mean across all FC values) as quality estimator but the reported correlation with motion was not statistically significant.

Other methods entail identification and exclusion of time points for which head movement exceeds a certain threshold (Power et al., 2014; Patel et al., 2014). Such threshold becomes increasingly stringent as the effects of motion have received greater recognition (Engelhardt et al., 2017). We did not investigate such measures (e.g. Δ*r* reported in several articles (Power et al., 2012, 2013, 2014, 2019)) because it requires data scrubbing and our goal was to avoid discarding any frames/time points.

Corrections of group-level statistics have been already implemented by regressing a summary motion metric for each subject (Satterthwaite et al., 2012; Van Dijk et al., 2012; Yan et al., 2013a; Power et al., 2014) but we propose that adding TFC measure could bring further advantages.

### 4.2. Effect of ROI size

The impossibility of optimal brain MRI parcellation makes the definition of regions of interest arbitrary. The number of ROIs ranges from 10 to 10^4^ in voxel-based studies (for review see Zalesky et al., 2010; Shen et al., 2013). However, how ROI size affects functional connectivity is unclear, therefore we examined the quality of FC matrices of varying sizes with respect to motion; size of FC matrices varied from 10 to 840 ROIs according to Craddock atlas.

We found an effect of ROI size on the FC quality, i.e. finer parcellation yielded noisier FC matrices. According to QC-FC values this effect is not related to head movements as median QC-FC and QC-FC correlation with distance were constant across atlases. Using TFC confirmed that the decrease in quality is specifically related to other types of noise, only very large ROIs (atlas with < 100 ROIs) showed increasing absolute correlation between TFC and DVARS/FD with decreasing ROI size. However, very a large ROI carry the risk that the mean time course of the ROI may not represent any of the constituent time courses if different functional areas are included within a single ROI (Shen et al., 2013). Moreover, as we show if analyzing too few regions it is more difficult to establish a reliable relationship with gross head motion.

Using tSNR we analyzed if ROI size also affects BOLD signals quality. tSNR is a well-established estimator of data quality, considering all types of noise. Unfortunately, the tSNR value is highly dependent on recording parameters and thus it is difficult to compare it across studies. Nevertheless, similarly to Van Dijk et al. (2012), who reported strong Pearson correlation between voxel-based tSNR and RMS (*r*_*P*_ = 0.57, *p* < 0.001), we also obtained significant Spearman correlations between tSNR and both mean FD (*r*_*S*_ = −0.35, *p* < 10^−7^) and mean DVARS (*r*_*S*_ = −0.68, *p* < 10^−16^) for the most detailed Craddock atlas that corresponds the most to the voxel-based tSNR. According to Fig.4 BOLD signal of more detailed atlases is noisier compared to less detailed atlases. This effect was specifically related only to DVARS. Such results suggest that there is an increasing effect of noise on the BOLD signal.

In conclusion both time-series and FC matrices based on smaller ROIs are noisier and it is the size of regions (number of voxels) and not the number of regions that plays the key role here. Moreover, we argue that motion is not the main driving effect behind this quality decrease. In all fMRI studies it is advised that applied atlas parcellation should be chosen carefully with respect to the application and expected outcomes. Our finding that the less detailed FC matrices are of better quality is useful for all functional connectivity studies when detailed FC matrix is not necessary, so finer brain parcellation can be sacrificed for more robust estimates of connectivity. Our recommendation here is in line with the one of Zalesky et al. (2010) that if possible, less detailed atlases will produce more robust results because they are less susceptible to noise. Nevertheless, large ROIs must be created carefully, and we do not recommend using Craddock atlas with less than 100 ROIs.

### 4.3. Limitations and future directions

To ensure robustness of our findings, we have replicated the analysis on the HCP dataset. We analyzed preprocessed rs-fMRI of 1081 subjects (Van Essen et al., 2013; Smith et al., 2013). HCP dataset benefits from very low temporal resolution (Ugurbil et al., 2013) with TR=0.72 s per volume. HCP data have undergone several denoising techniques designed to remove artefacts (HCP FIX-ICA denoising, motion regression and censoring high-motion time points) with the intention of providing clean data (Marcus et al., 2013; Burgess et al., 2016). We replicated all our obtained results and proved TFC to be the most reliable FC quality estimator (Fig.F.9). The obtained correlations were generally lower, but this might be explained by more strict preprocessing (including censoring time points). That is the reason why the QC-FC correlation diminished as also reported in Ciric et al. (2017) where ICA-AROMA was the only method to show virtually no QC-FC distance-dependence. Again, we did not find a significant change in the TFC-motion relationship except for the very small atlases. However, we observed a progression of the tSNR-motion relationship for both DVARS and FD (Fig.F.8).

The question arises as to which motion metric is optimal. Currently, the most used motion parameters across studies are DVARS and FD (Waheed et al., 2016). As Power et al. (2012) pointed out it is difficult to quantify the effect of motion with only one parameter. Nevertheless, according to our results mean DVARS is strongly correlated with FC quality (*r*_*S*_ up to 0.5). Other summarizing parameters such as maximum of DVARS or variance of DVARS could be used as well because they capture other features of motion (big spike-like movements, constant small drift) but Van Dijk et al. (2012) showed that they are all highly correlated (i.e. mean motion was strongly correlated with both max motion and a number of movements) therefore we only reported mean FD and mean DVARS. Naturally, the accuracy of techniques that use FD and DVARS are limited by the accuracy of the measures themselves (Power et al., 2015). Since motion takes the form of regionally heterogeneous effects on functional connectivity estimates, better measurements of motion can yield better predictions for FC quality. For example, a shorter TR has the effect of dividing large movements into several smaller movements. That is why rapid sub-TR displacements were thought to play a significant role in regional motion artefact interactions (Spisak et al., 2014). Nevertheless, previous studies found that sub-TR FD traces are noisier and less useful in identifying outlying time points, but DVARS traces exhibit signal-to-noise ratios that are useful for identifying outlying data points (Power et al., 2014). Therefore, sub-TR DVARS metric could be more accurate estimate of subject’s movement. Nonetheless, we did not observe any improvement in the HCP dataset with lower temporal resolution (TR=0.72 s). Another possible improvement could be to use slice-derived motion metrics rather than volume-derived estimates because they are only a simplification of movement over the acquisition of all slices (Beall and Lowe, 2014). Anyway, Satterthwaite et al. (2013) and Yan et al. (2013a) found that motion correction with voxel-wise motion metrics offered insufficient advantage over the more easily computed general models.

Unfortunately, we are not able to provide a single value that would separate bad and good FC matrix due to the complexity of all contributing factors, such as the lack of a ground truth of FC. Therefore, the decision on which scanning session should be discarded is still based only on a summary motion statistic reaching some threshold (i.e. RMS movement over half a voxels width (Power et al., 2013) or more than 20 volumes with RMS greater than 0.25 mm (Ciric et al., 2017)). We only propose to add TFC measure for group-level corrections. Other directions of mitigating the motion artefact include using multi-echo imaging (Power et al., 2018) or using head molds (Power et al., 2019).

A possible objection is that the typical connectivity matrix is not an appropriate golden standard. Nonetheless, we assume that by averaging FC matrices of low-movement subjects we obtain a typical awake human brain functional connectivity free of motion artefact (as well as other biases). We also found that the group-average FC matrices from different groups were very similar (correlation of the typical matrix with similarly preprocessed typical FC of the different dataset is *r*_*P*_ = 0.91, reps. *r*_*P*_ = 0.68 with HCP dataset), therefore we obtained similar results regardless of what typical FC matrix was used. Moreover, using the typical FC matrix from a different dataset has the advantage that no degrees of freedom are lost, i.e. subjects used for the computation of the typical FC matrix do not have to be discarded from subsequent analyses.

## 5. Conclusion

In this paper we present a new method of functional connectivity quality evaluation for rs-fMRI data. Typicality of Functional Connectivity metric is based on a correlation of subjects FC matrix with the low-motion group-average FC matrix. This metric is strongly correlated with motion metrics and it allows for the assessment of the effect of head movement on individual connectivity matrices. Therefore, TFC provides an individual proxy measure of motion effect on functional connectivity and is more sensitive to inter-individual variation of motion than individual functional connections. We used it to show that there is a gradual decrease of the connectivity quality and the data quality in more detailed atlases with ROIs composed of fewer voxels. Nevertheless, the motion connectivity quality relationship remained stable across atlases because the effect of motion does not depend on the atlas size unlike other possible types of noise. Such knowledge is useful in several scenarios such as group comparison, preprocessing pipeline performance estimation and choosing brain parcellation. Moreover, outcomes of different analyses can vary significantly according to the brain parcellation employed. These findings should be considered when a robust estimate of connectivity is more important than fine brain parcellation or while comparing denoising strategies.

## Acknowledgement

This work has been supported by the Czech Science Foundation projects No. 13-23940S and 17-01251S, project Nr. LO1611 with a financial support from the MEYS under the NPU I program.

## Appendix A. Alternative dataset

84 healthy controls (80 right-handed, 48 males, mean age 30.83 / standard deviation 8.48) participated in the control study. Participants were informed about the experimental procedures and provided written informed consent. The study design was approved by the local Ethics Committee of the Institute of Clinical and Experimental Medicine and the Psychiatric Center Prague. Each volunteer underwent MRI scanning that included 10 minutes of resting-state functional magnetic resonance imaging acquisitions with eyes closed and an acquisition of a T1-weighted and T2-weighted anatomical scan.

Scanning was performed with a 3T MR scanner (Siemens; Magnetom Trio) located at the Institute of Clinical and Experimental Medicine in Prague, Czech Republic. Functional images were obtained using T2-weighted echo-planar imaging (EPI) with blood oxygenation level-dependent (BOLD) contrast. GE-EPIs (TR/TE=2500/30 ms) comprised axial slices acquired continuously in descending order covering the entire cerebrum (voxel size=2 × 2 × 2mm^3^). A three-dimensional high-resolution T1-weighted image (TR/TE/TI = 2300/4.6/900 ms, voxel =1 × 1 × 1mm^3^) covering the entire brain was used for anatomical reference. T2-weighted images were also acquired, but not used in the current study.

## Appendix B. Preprocessing

### Appendix B.1. Moderate

In comparison with stringent pipeline the moderate denoising steps only included regression of six head-motion parameters (acquired while performing the correction of head-motion) and one principal components of white-matter and cerebrospinal fluid. A band-pass filter with broader cutoff frequencies, i.e. 0.004 - 0.1 Hz, was applied.

### Appendix B.2. Raw

Raw preprocessing consists of only CONNs default preprocessing pipeline (defaultMNI): (1) functional realignment and unwarping, (2) slice-timing correction, (3) structural segmentation into white matter and cerebrospinal fluid & structural normalization to the MNI space, (4) functional normalization to the MNI space, (5) outlier detection, and (6) smoothing with 8mm kernel size. No further filtering or regression was done.

## Appendix C. Dataset motion parameters

Mean of DVARS and FD are metrics are commonly used to described gross head movement of a given subject. In Fig.C.6. we provide a distribution of such metrics for the 245 subjects used in this study.

**Figure C.6:**
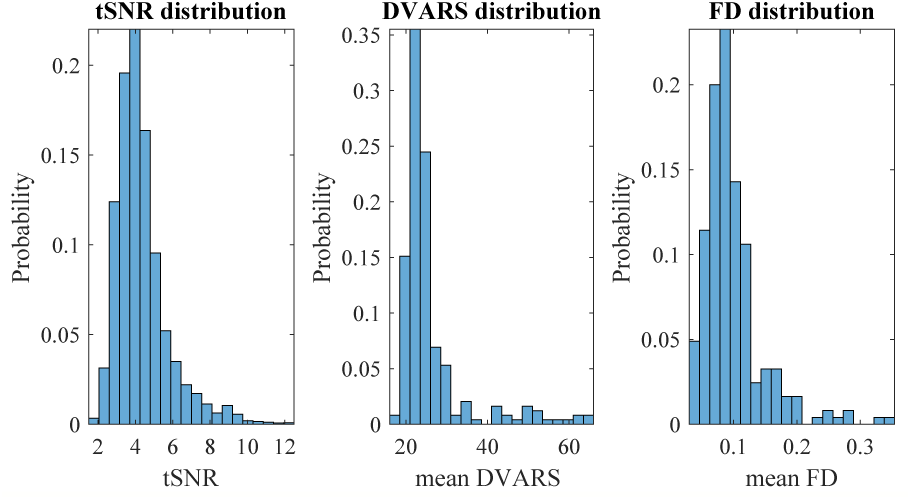
Distribution of mean FD and mean DVARS in the dataset. All subjects would meet inclusion criteria for analysis in most MRI laboratories.

## Appendix D. Examples of FC matrices

Subjects with low FCT have FC matrices less resembling typical FC matrix. According to Fig.D.7 this degradation is apparent during a visual inspection.

## Appendix E. tSNR

Temporal signal to noise ratio is a useful measure of data quality (Bodurka et al., 2007). Van Dijk et al. (2012) have found that low values of tSNR identify subjects with high head motion or other causes of data instability. For each ROI the mean signal is divided by the standard deviation over the BOLD run. Then, tSNR is the mean tSNR value across all ROIs in the brain (Eq.E.1).

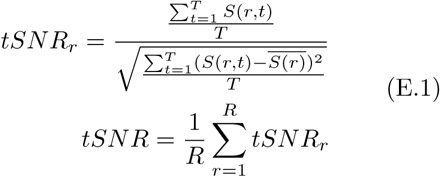

where *S*(*r, t*) is the signal magnitude at the ROI *r* in the time *t*. 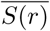 is a temporal mean. *T* is number of all brain volumes and *R* is number of all ROIs.

## Appendix F. Human Connectome Project

We analyzed preprocessed rs-fMRI of 1081 subjects (Van Essen et al., 2013; Smith et al., 2013) and performed the same analyzes as described in the Methods and Results section.

**Figure D.7:**
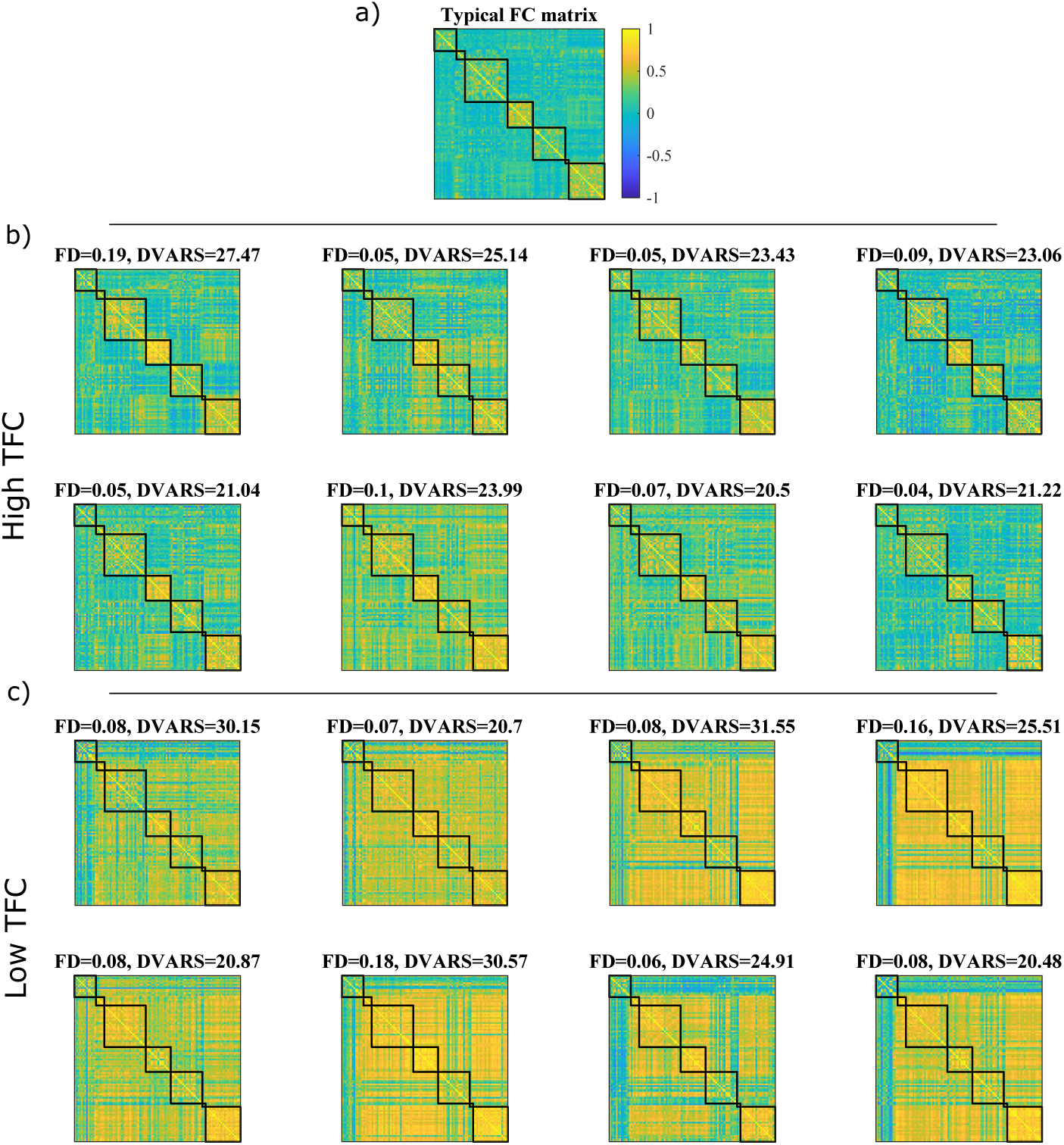
Examples of FC matrices with moderate preprocessing for different subjects based on their TFC. a) Typical FC matrix for Craddock atlas with 100 ROIs. b) Examples of subjects with lowest TFC. Based on visual inspection, their FC matrices do not correspond to the typical matrix, even though their head movement might not be prominent. c) Examples of subjects with highest TFC. Their FC matrix nicely preserved the modular structure similar to the typical matrix.

**Figure F.8:**
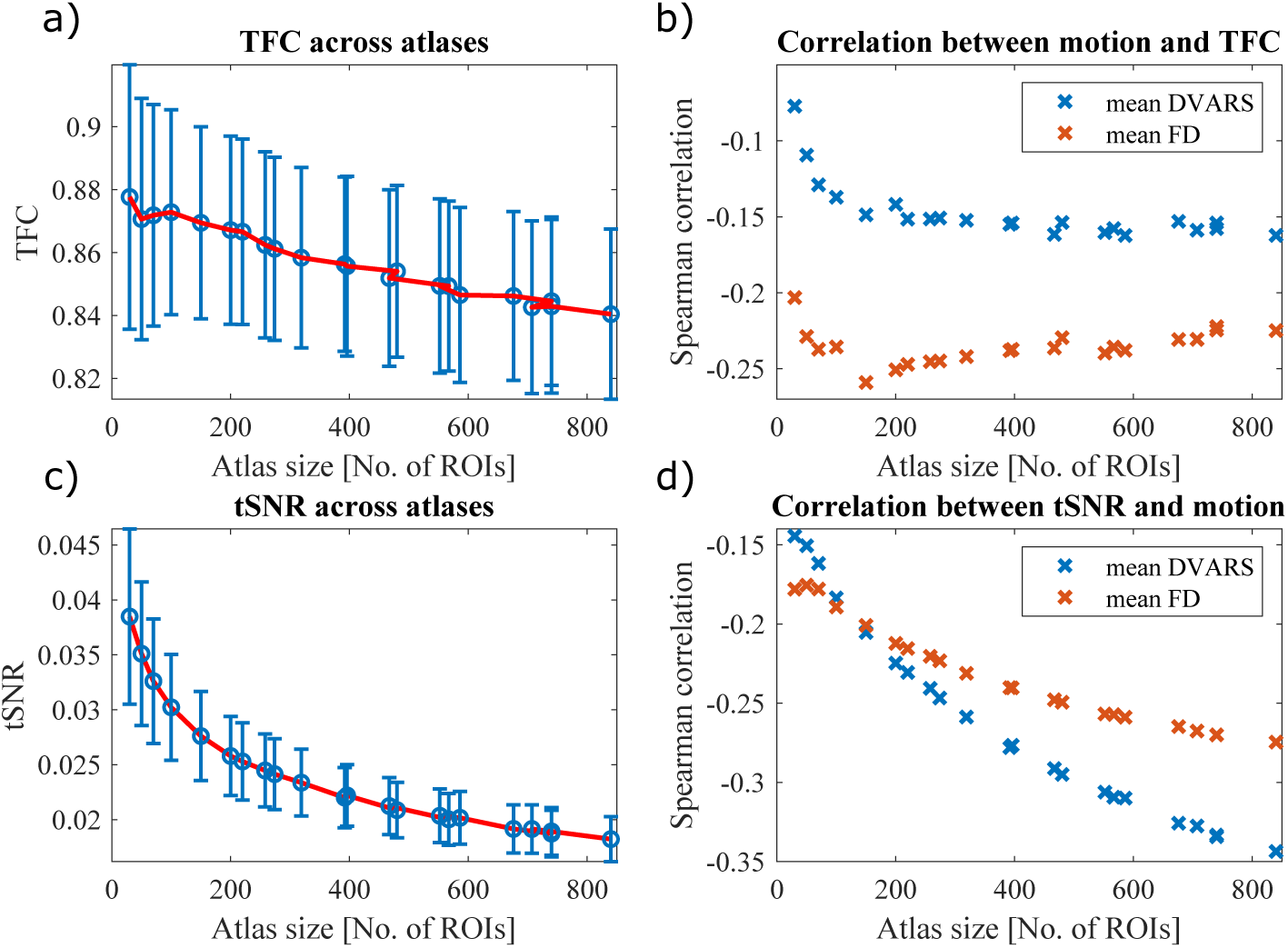
HCP dataset quality assessment. a) Dependence of data quality on atlas size. TFC magnitudes are comparable to those obtained in our datasets. Similarly, TFC is decreasing with decreasing atlas size. b) We confirm that the TFC-motion relationship is stable across various atlases (except for the smallest ones) and that DVARS shows stronger absolute correlation with TFC. c) Decreasing atlas size mean also decreasing tSNR in HCP dataset. d) The gradually increasing absolute correlation between tSNR and DVARS is still present and, unlike in our dataset, there is also a graduate increase in the tSNR-FD correlation. Therefore, there is an effect of atlas size in HCP dataset as well.

**Figure F.9:**
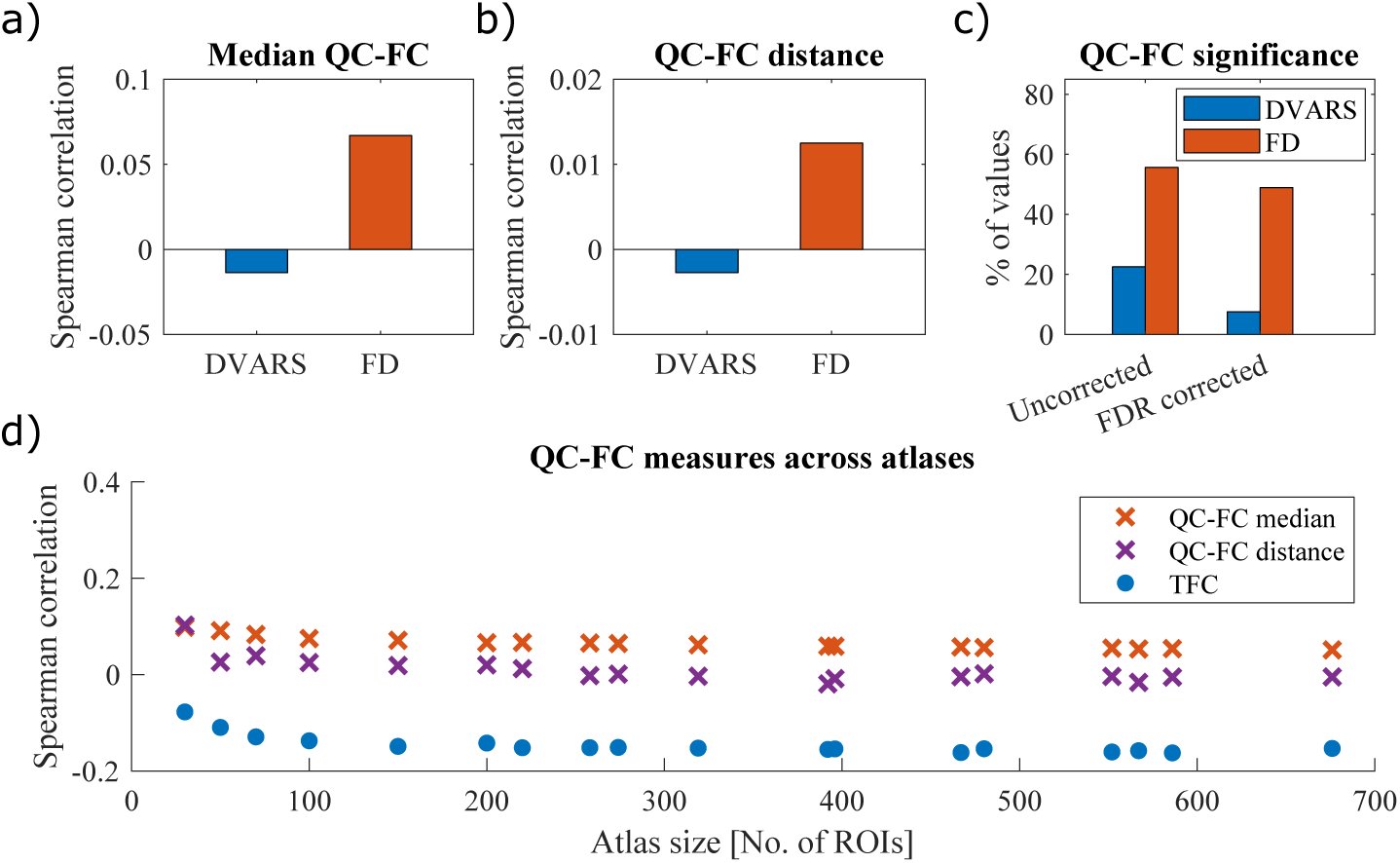
Estimation of the effect of head motion present in HCP dataset using QC-FC values. a) Only the median FD-FC values shows a spurious increase of connectivity b) Moreover, we did not obtain a significant correlation between QC-FC values and distance proving a successful mitigation of distance dependence and other motion-related impurities for the HCP preprocessing pipeline. c) Nevertheless, QC-FC relatively high amount of FC values is correlated with head movements (> 50 % for FD). d) Still, TFC was able to relate the motion (mean FD) to FC quality proving its usefulness as an estimator of studied relationships. Based both on QC-FC and TFC the head motion effect on connectivity seems to be constant and independent of ROI size.

